# Intracellular Vesicle Entrapment of Nanobubble Ultrasound Contrast Agents Targeted to PSMA Promotes Prolonged Enhancement and Stability *In Vivo* and *In Vitro*

**DOI:** 10.1101/2021.05.11.443511

**Authors:** Reshani H. Perera, Eric Abenojar, Pinunta Nittayacharn, Xinning Wang, Gopal Ramamurthy, Pubudu Peiris, Ilya Bederman, James P. Basilion, Agata A. Exner

## Abstract

Previous work has shown that active targeting of nanobubble (NB) ultrasound contrast agents to the prostate-specific membrane antigen (PSMA) significantly prolongs ultrasound signal enhancement in PSMA-expressing prostate cancer. However, the specific mechanism behind this effect is not well understood. Furthermore, prior studies were carried out using clinical ultrasound scanners in a single imaging plane. Because tumor heterogeneity can have a drastic effect on bubble kinetics and resulting contrast enhancement, a single region of interest in one imaging plane over time may not fully represent the contrast dynamics of the entire tumor. Accordingly, in the current work, we used high-frequency dynamic parametric contrast-enhanced ultrasound (DCE-US) imaging to gain a detailed understanding of NB kinetics in prostate tumors in mice. Specifically, we examined the differences in enhancement between the tumor periphery and tumor core in the same imaging plane. We also quantified intact nanobubble retention in the entire tumor volume. To better understand the mechanism behind prolonged tumor enhancement, intracellular retention and the acoustic activity of PSMA-NB were evaluated in cell culture. DCE-US US data suggest that both tumor wash-in and retention of PSMA-NB are delayed due to biomarker interaction and binding. The longer retention of PSMA-NB signal in tumor core supported target-driven bubble extravasation. *In vitro* studies demonstrated a higher level of internalization and prolonged-acoustic activity of internalized PSMA-NB. GC/MS analysis confirmed gas persistence in the cells after PSMA-NB internalization. The active-targeting of NB results in cellular internalization via receptor-mediated endocytosis, and the location with intracellular vesicles (late-stage endosomes/lysosomes) significantly prolongs gas retention within the cells. These features can enable background-free diagnostic imaging of the target cells/tissues, as well as highly focused ultrasound-modulated therapeutic interventions.

## 1. Introduction

Prostate specific membrane antigen (PSMA) is a well-established biomarker for prostate cancer (PCa).^[1–3]^ PSMA is a membrane bound glycoprotein that is highly expressed on the cell surface of PCa cells. Previously, we reported on the development of nanobubble (NB) ultrasound (US) contrast agent that specifically targets the PSMA receptor in PCa^[4, 5]^ via the targeting moiety PSMA-1. PSMA-1 is a urea-based ligand that has a high affinity to PSMA (IC_50_=2.01nM)^[6, 7]^. The PSMA-1 targeted NB (PSMA-NB) previously showed selective uptake into PSMA expressing PC3pip cells and a significantly higher accumulation in PSMA-positive tumors ^[4, 5]^. However, in light of significant tumor heterogeneity, data from a single imaging plane does not fully capture the contrast dynamics in the entire volume. Due to the high proliferative rate of cancer cells, blood vessels develop rapidly and form an immature vascular network that contributes to the vascular heterogeneity throughout the same lesion and among tumors^[8],[9]^. Variations in microvessel density and permeability can have a direct effect on nanobubble extravasation into the tumor parenchyma. Likewise, variation in the tumor microenvironment, such as cancer cell density, presence of immune cells, fibrosis and interstitial pressure, can all affect nanobubble distribution and retention within the tumor and uptake into tumor cells. All of these factors will affect the apparent NB contrast agent dynamics which are captured on the time intensity curves (TIC), but these may not be accurately represented by average region of interest (ROI) data analysis over the entire tumor area. The current study thus aimed to investigate the kinetics of PSMA-targeted NB distribution with high frequency ultrasound across the entire tumor volume, and sought to examine the differences in contrast agent in the tumor rim and tumor core. The work also further validates the PSMA-NB extravasation and accumulation in whole tumor mass using three dimensional (3D) US imaging.

We previously also showed that active targeting to PSMA enhances tumor uptake of *intact* PSMA-NBs with extended retention, which results in the prolonged US signal enhancement in the tumors that can be visualized with clinical nonlinear ultrasound.^[4]^ One hypothesis for the prolonged tumor enhancement is that PSMA-targeted NBs are internalized into their target cancer cells and the internalization delays octafluoropropane gas dissolution. The interaction of microbubbles (MBs) and cells has been reported in several studies, showing that MBs can enter the cell during sonoporation, associate with the plasma membrane, or can attach to cell surface receptors when targeted. Some of these interactions have been shown to slow the decay of ultrasound signal generated from the MBs.^[10–16]^ Additional studies have reported on the effects of MB stability and inertial cavitation on cellular trafficking of drugs and dyes.^[12]^ However, no work has specifically examined and validated the kinetics of cellular internalization of NBs and compared these processes in the case of free and targeted nanobubbles. Thus, this study also aimed to investigate the effect of receptor-mediated endocytosis of PSMA-NBs on their acoustic activity and intracellular persistence using an *in vitro* cellular model. Elucidating the mechanism of interaction of PSMA-NB at the cellular level can offer a new avenue to clinical translation of PSMA-NB in PCa diagnosis and therapeutic applications.

## 2. Experimental section/Methods

### 2.1 NB formulation and characterization

The preparation and characterization of NBs has been reported elsewhere.^[17–19]^ Briefly, a cocktail of lipids including DBPC (Avanti Polar Lipids Inc., Pelham, AL), DPPE, DPPA (Corden Pharma, Switzerland), and mPEG-DSPE2000 (Laysan Lipids, Arab, AL) were dissolved in propylene glycol (PG, Sigma Aldrich, Milwaukee, WI), glycerol and PBS. Then gas exchanged with C_3_F_8_ (Electronic Fluorocarbons, LLC, PA) and vial was subjected to mechanical agitation. NBs were isolated by centrifugation. PSMA targeted NB formulated by incorporating DSPE-PEG-PSMA-1 into the lipid cocktail mixture^12^. PSMA-NB and NB were characterized with resonant mass measurement (Archimedes®, Malvern Panalytical) and validated using HPLC and Matrix-assisted laser desorption/ionization time-of-flight mass spectroscopy (MALDI-TOF-MS) as previously described.^[5, 18]^

### 2.2 Cell culture studies

Retrovirally transformed PSMA-positive PC3pip cells and PC3flu cells (transfection-control) were originally obtained from Dr. Michel Sadelain (Memorial-Sloan Kettering Cancer Center, New York, NY).^[20]^ Cell lines were checked and authenticated by western blot. Cells were grown in complete RPMI1640 medium (Invitrogen Life Technology, Grand Island, NY) at 37 ° C and 5% CO_2_ environment.

### 2.3 Animal model

Animals were handled according to a protocol approved by the Institutional Animal Care and Use Committee (IACUC) at Case Western Reserve University and were in accordance with all applicable protocols and guidelines in regards to animal use. Male athymic nude mice (4-6 weeks old) were anesthetized with inhalation of 3% isoflurane with 1L/min oxygen and were implanted subcutaneously with 1×10^6^ of PSMA-positive-PC3pip cells in 100 μL Matrigel. Animals were observed every other day until tumors reached at about 8-10 mm in diameter.

### 2.4 In vivo NLC imaging for bubble kinetic analysis

*In vivo* experiments were performed using FUJIFILM VisualSonics Vevo 3100. A total of 9 animals were used for this experiment. Animals were divided into 3 groups; PSMA-NB, NB and Lumason MB group. *In vivo* bubble distribution was imaged with 2D nonlinear contrast mode. A total volume of 200 μl of undiluted either PSMA-NB or NB were injected via tail vein. To obtain wash-in bubble dynamic tumor scanned for 3.33min at 5 fps. The scanning parameters were setup to 18 MHz frequency, with MS250 transducer, 4% transmit power, 30 dB contrast gain, medium beam width, 40 dB dynamic range. Then 3D US scan was then performed at the peak signal. Transducer was clipped onto the 3D motor and positioned onto the tumor area. By adjusting the X-Y axis position of the probe, placed the probe at the center of the tumor as the imaging display. The 3D setup arranged to make 0.05 mm size thickness 2D slices with 383 frames. Images obtained and constructed together to obtain the 3D volume to achieve whole tumor bubble distribution. Then non-linear contrast imaging was accomplished to see the kinetic of washout phase at 1fps and 1000 frames for ~ 16 min with maintaining above parameters during the imaging session. After NLC, 3D imaging was performed. To confirm intact bubbles extravasated and accumulated in tumor 3D burst sequence was applied to entire tumor^[21]^ and rescanned to obtain 3D image.

### 2.5 Tumor extravasation studies with 3D ultrasound

For the extravasation studies, animals were divided into 3 groups: PSMA-NB, NB and Lumason, and 3 animals was used for each group (total n = 9). 200 μl of contrast agent was injected via tail vein. Mouse was subjected to 3D US scan 25 min post injection as described as above. Then cardiac perfusion was performed with 50 ml of PBS through the left ventricle and 3D US scan was completed again to detect the US signal produced from intact bubbles that accumulated in the perfused tumor.

### 2.6 Histology analysis

Animals were divided into three groups: Cy5.5-PSMA-NB (n =3), Cy5.5-NB (n=3), and no-contrast-control. Cy5.5 labeled NBs were prepared by mixing DSPE-PEG-Cy5.5 (100μl) into the lipid solution. Mice received either 200μl of undiluted UCAs or PBS via tail-vein. 25 min after injection, animals were scanned using US to detect the signal and then PBS perfusion was performed with 50ml-PBS though left-ventricle. Then, tumors were scan again to perceive the US signal that generate from intact-NB. Tumors and the kidney were harvested, fixed in paraformaldehyde and embedded in optimal-cutting-temperature compound (OCT Sakura Finetek USA Inc., Torrance, CA). The tissues were cut into 8μm slices and washed (3X) with PBS and incubate with protein blocking solution that contain 0.5% TritonX-100 (Fisher Scientific, Hampton, NH) and incubated in 1:250 diluted primary-antibody CD31(PECAM-1) Monoclonal Antibody Fisher Scientific, Hampton, NH) for 24h at 4°C. It was then washed with PBS, incubated with Alexa-568 tagged secondary-antibody (Fisher Scientific, Hampton, NH) for 1h, and stained with DAPI (Vector Laboratories, Burlingame, CA). The fluorescence images were obtained and analyzed (by interactive function of segmentation and threshold) using Axio Vision V 4.8.1.0, Carl Zeiss software (Thornwood, NY). For PSMA-immunohistochemistry, tissues were wash 3X with PBS and incubated with blocking solution followed by 1:150 diluted PSMA primary-antibody (Thermo Fisher Scientific, Waltham, MA) for 24h at 4°C and followed the above steps as for CD31 staining.

### 2.7 Confocal imaging of PSMA-NB internalized PC3pip cells

PC3pip cells were seeded in glass bottom petri dishes (MetTek Corporation, Ashland, MA, USA) at a density of 10^4^ cells/well. Rhodamine-labeled-NBs were prepared by mixing DSPE-Rhodamine (50 μl) into the lipid solution. After 24 h, 1: 30 diluted Rhodamine-tagged PSMA-NB (250 μL) were added to the cells for 1h. Following incubation, cells were washed with PBS and were then placed into the incubator in RPMI for 3 h and for 24 h. The 3 h time-point was chosen because a significantly high acoustic activity was previously observed with PSMA-NB internalized PC3pip cells with low standard error at the 3 h time point. One hour before the end of incubation 5 μM Lysotracker Red 24 μL), a marker for late endosomes and lysosomes, (ThermoFisher Scientific) was added the cells, as per manufacturer instructions. After incubation, cells were washed 3X with PBS and fixed with 4% paraformaldehyde for 10 min. Cells were washed with PBS and stained with DAPI mounting medium (Vecor Laboratories, Burlingame, CA). Then cells were observed using a fluorescent microscope (Leica DMI 4000B, Wetzlar, Germany) equipped with the appropriate filter sets. LysoTracker Red exhibits green fluorescence (excitation: 577 nm, emission: 590 nm).

### 2.8 Cellular uptake studies

Both PC3pip and PC3flu cells (2 ×10^6^ cells/ml) were plated on cell culture petri dishes (60 × 15 mm, Fisher Scientific) at about 70 % confluence. Twenty-four hours later, cells were incubated with PSMA-NB or plain NBs (~ 10,000 bubbles/cell) for 1 h. After incubation, cells were washed three times with PBS (3X) and maintained in RPMI at 37 ° C until the US scan time points. Before US scan cells were trypsinized, counted and 1 ×10^6^ cells were used.

### 2.9 Acoustic assessment of PSMA targeted NB internalized in PC3pip cells *in vitro*

*In vitro* acoustic activity of NBs internalized cells was assessed using a clinical US scanner (AplioXG SSA-790A, Toshiba Medical, now Hitachi Healthcare America). To carry out the measurements, cells (~ 2× 10^6^) were washed and detached using trypsin. Following detachment and resuspension in PBS, the cell suspension was placed in a custom-made 1.5% (w/v) agarose phantom.^[19]^ The phantom was affixed over a 12 MHz linear array transducer, and images were acquired with contrast harmonic imaging (CHI) at 0.1 mechanical index (MI), 65 dynamic range, 70 dB gain, and 0.2 frames/sec imaging frame rate. Using onboard software, a region of interest (ROI) analysis was performed on all samples to measure the mean signal intensity in each ROI.^[22]^ The data were then exported to Microsoft Excel for further processing. The experiments were carried out in triplicate.

### 2.10 Analysis of octafluoropropane gas trapped in cells using GC/MS

Presence of octafluoropropane (C_3_F_8_) gas inside cells was confirmed by headspace gas chromatography / mass spectrometry (GC/MS) as previously described.^[18]^ For these experiments, cells (1×10^7^ cell/mL) were grown in cell culture flasks (75 cm^2^ size) for 24 h. Again, as above, cells were then incubated with 1 mL of either PSMA-NB or plain NB (~ 10000 NBs/cell) for 1 h. After incubation, cells were washed with PBS and incubated in medium for 3 h. Following incubation, the cells were trypsinized, re-suspended in PBS and centrifuged at 1000 rpm for 4 min. Then the cells were then transferred to GC headspace vials with 300 μL of medium and 300 μL of cell lysis buffer, sealed with PTFE/silicon septum and capped (Thermos Fisher Scientific). The vials were sonicated for 20 min in an ultrasonic water bath (Branson Ultrasonics, Danbury, CT) at 50 ° C to release C_3_F_8_ gas into the headspace vial. The GC/MS analysis was performed as described previously^[18]^ using the Agilent 5977B-MSD equipped mass spectrometer with an Agilent 7890B gas chromatograph GC/MS system. A DB5-MS capillary column (30 m × 0.25 mm × 0.25 μm) was used with a helium flow of 1.5 mL/min. Headspace samples of 1 μL were injected at 1:10 split. Gas chromatography conditions used were as follows: oven temperature was at 60 °C, held for 1 min, ramp 40 °C/min until 120 °C and held for 3.5 min. Perfluoropropane eluted at 1.2 min. Samples were analyzed in Selected Ion Monitoring (SIM) mode using electron impact ionization (EI). M/z of 169 (M-19) was used in all analyses. Ion dwell time was set to 10 ms. Perfluoropropane was verified by NIST MS spectra database. The standard calibration plot was obtained by measuring the peak area of GC peak as a function of NB concentrations (0-100 × 10^8^ NB/mL).

### 2.11 *In vivo US* Imaging of internalized bubbles

Animals were handled according to the protocol approved by the Institutional Animal Care and Use Committee (IACUC) at Case Western Reserve University and were in accordance with all applicable protocols and guidelines in regards to animal use. PC3pip cells were incubated with PSMA-NB and processed as described above. Mice (n=9, 4-6 weeks old Athymic (NCR nu/nu) mice with 20g weight) were randomly divided into 3 groups and were anesthetized with inhalation of 3% isoflurane with 1 L/min oxygen. Baseline US signal was obtained both left and right side of flank area (marked with a permanent marker) using the parameters described above at 0.1 and 0.5 mechanical index (MI). Following incubation with PSMA-NB, the PC3pip cells were suspended in a Matrigel and PBS mixture (1:1 PBS/Matrigel) and cell suspension (100 μl) was injected subcutaneously into flank of nude mice. As a control, cells without NB exposure were injected adjacent to the NB-exposed cells (Figure 5A). US images of the injection sites were obtained immediately after injection at MI=0.1 immediately or after 3, 24 h or 8 days using the same parameters as above. After imaging at 0.1 MI, the MI value was increased to 0.5 and regions were imaged again for all time points. The region of interest (ROI) was drawn around the injected cell-region, excluding the skin. Then mean signal intensity for each ROI was obtained from the CHIQ software.^[22]^ These measurements were exported to Excel and the signal enhancement in each ROI by subtracting the baseline value (contrast before the inoculation).

**Figure 1.**
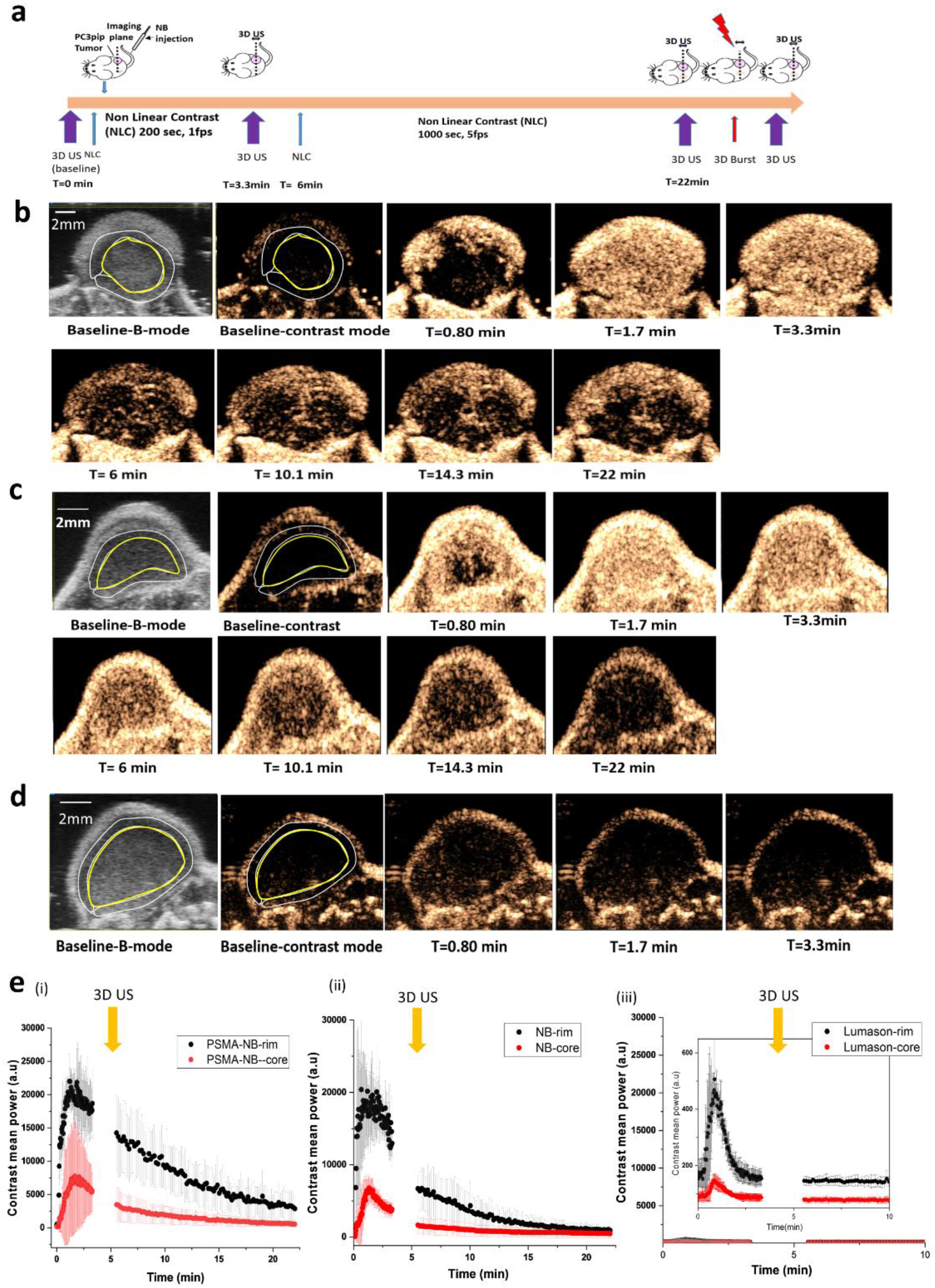
*In vivo* nonlinear contrast enhanced ultrasound of prostate tumors in mice. a) Timeline showing the experimental approach. b)-d) Representative US images showing tumor enhancement at different times following injection of PSMA-NB (b), Plain NB (c), and Lumason microbubbles (d). e) Average signal intensity in tumor rim and core as a function of time for PSMA-NB (i), plain NB (ii), and Lumason MB (iii). Tumors were imaged at 18 MHz, 5 frames/second for 3.33 min and 1 frame/s for 16.67 min.

**Figure 2.**
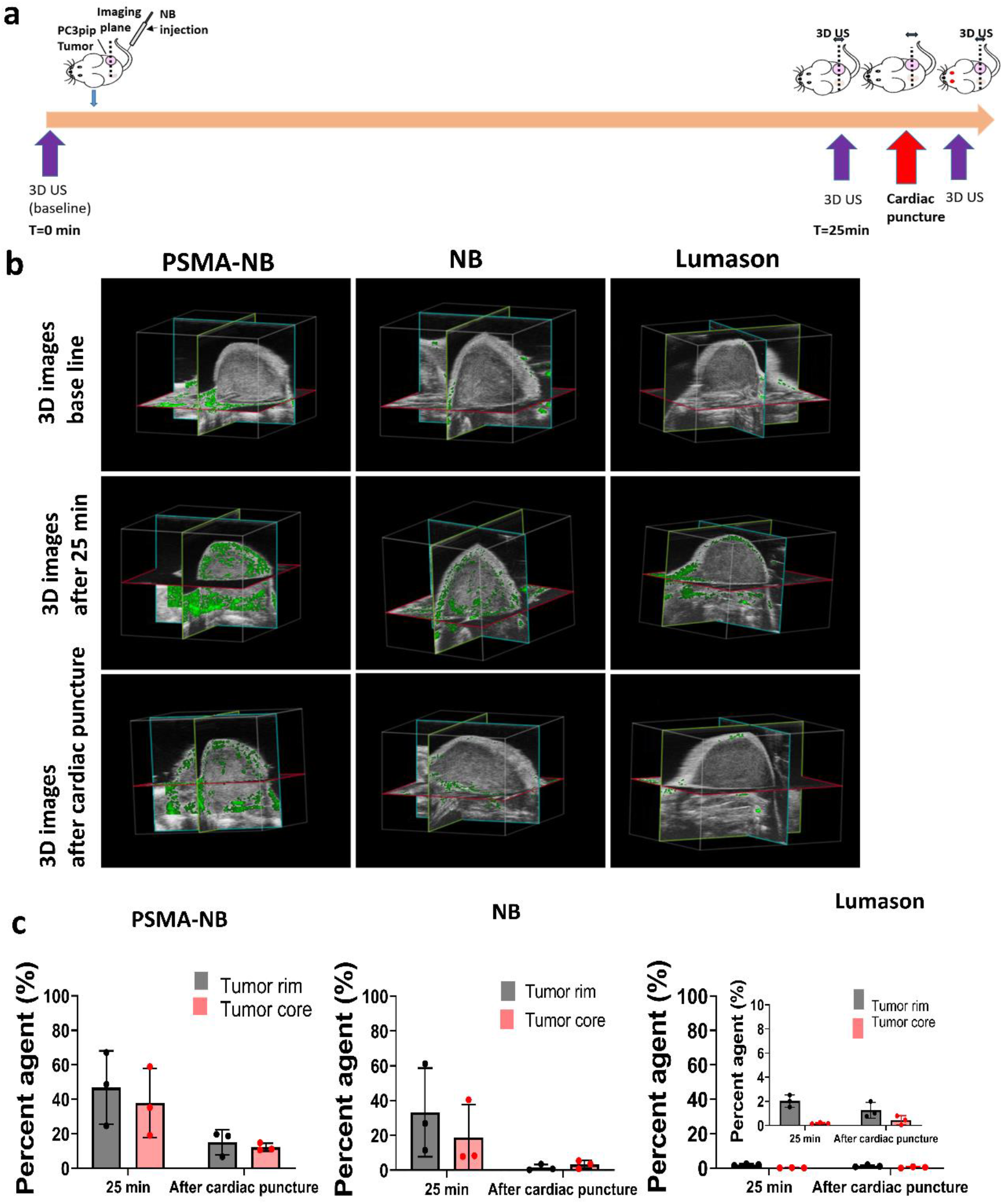
3D ultrasound scanning performed without constant exposure of tumors to US following NB injection. a) Timeline showing the 3D US scanning points. b) Representative 3D US images of the tumor showing nonlinear signal from PSMA-NB, NB, and Lumason at the baseline, 25 min post injection, and after cardiac puncture. c) Quantification of nonlinear signal in each tumor volume before and after cardiac puncture for PSMA-NB, plain NB, and Lumason after baseline subtraction. n=3, error bars represent mean ± s.d., * p<0.05.

**Figure 3.**
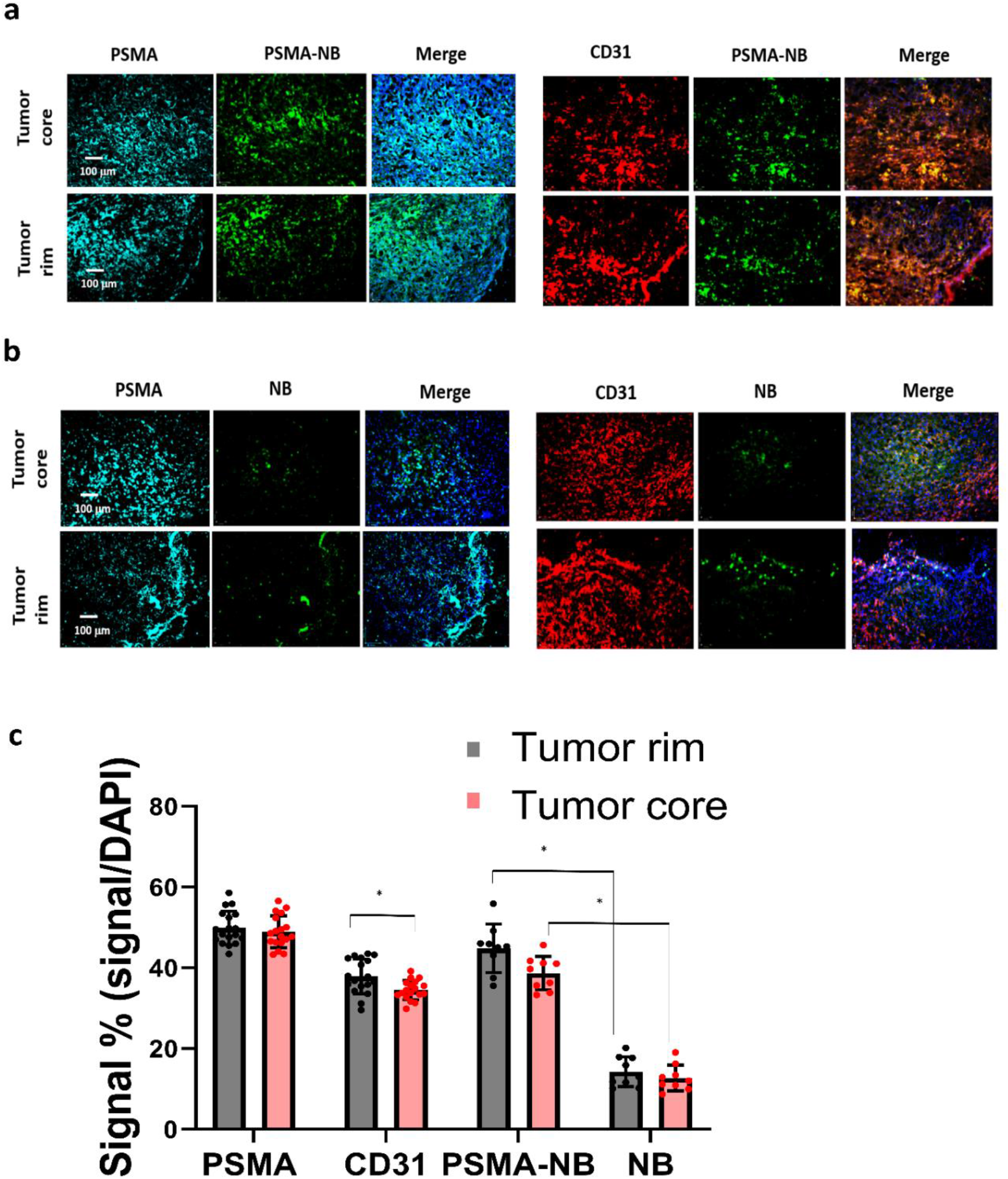
Histology images showing the Cy5.5-PSMA-NB accumulation and extravasation in PC3pip tumor that were excised after cardiac puncture. Representative images (10X magnification) of tumor tissues showing the PSMA expression (cyan), vasculature (CD31 expression, red), and distribution of a) PSMA-NB (green) or b) plain NB (green). c) The signal intensities of bubbles, PSMA and vessel are shown as the percentage of total DAPI (cell) fluorescence in tumor section. Cy5.5-PSMA-NB signal in both tumor rim and core was significantly higher from that of NB. n=3, error bars represent mean ± s.d., * p<0.001.

**Figure 4.**
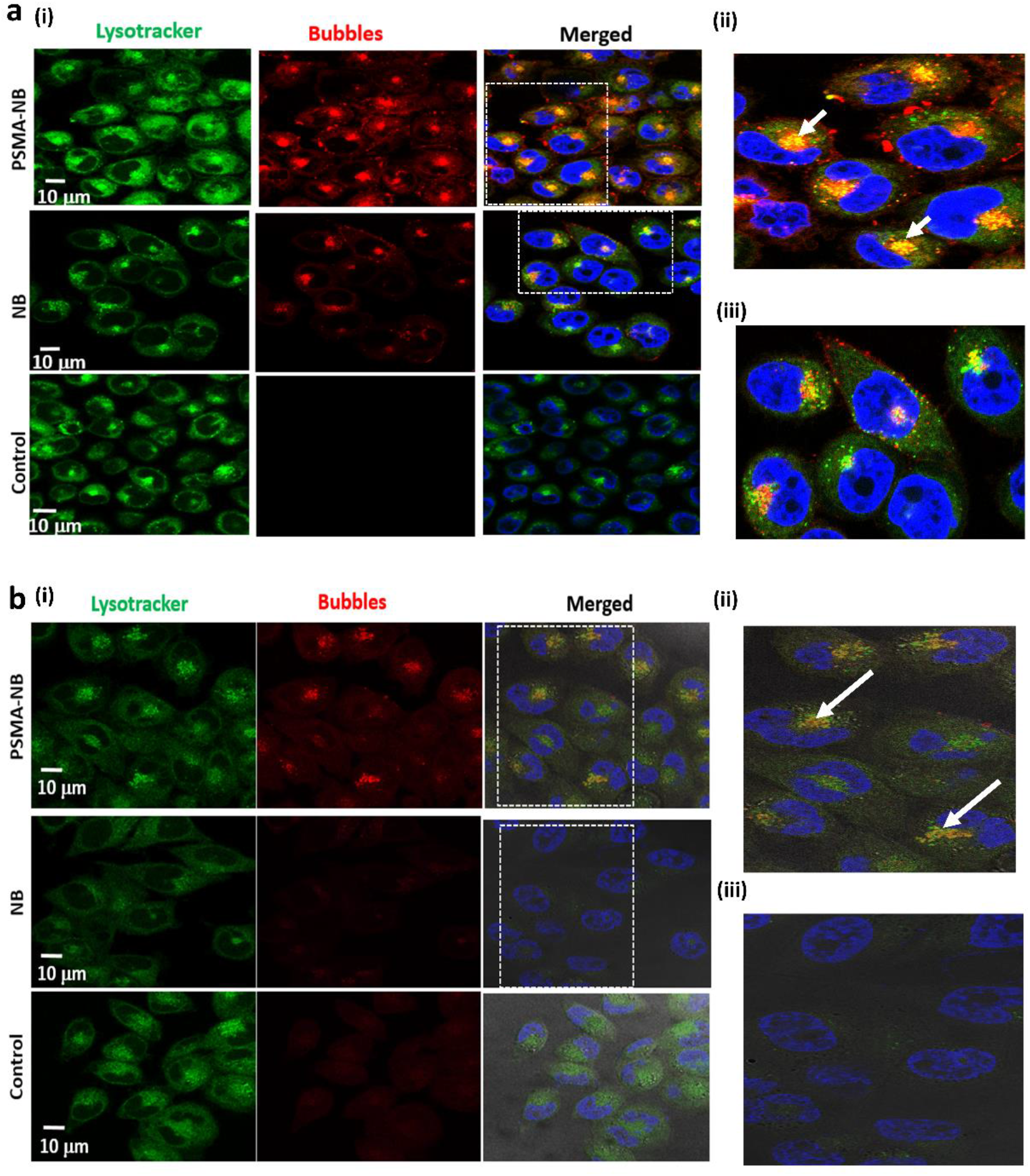
(a) Representative confocal images of PSMA-NB and NB distribution after 1h exposure in PC3pip cells; 100X (blue-nuclei, red-NB, and green-late endosome/ lysosomes). Zoomed merged images of (ii)PSMA-NB and (iii) plain NB incubated PC3pip cells. (b) Representative confocal images of PSMA-NB distribution in PC3pip cells after 24h exposure (blue-nuclei, red-NB, and green-endosome). Zoomed merged images of (ii) PSMA-NB and (iii) plain NB incubated PC3pip cells. PSMA-NB shows high co-localization in late endosomal/ lysosomal vesicles (yellow).

**Figure 5.**
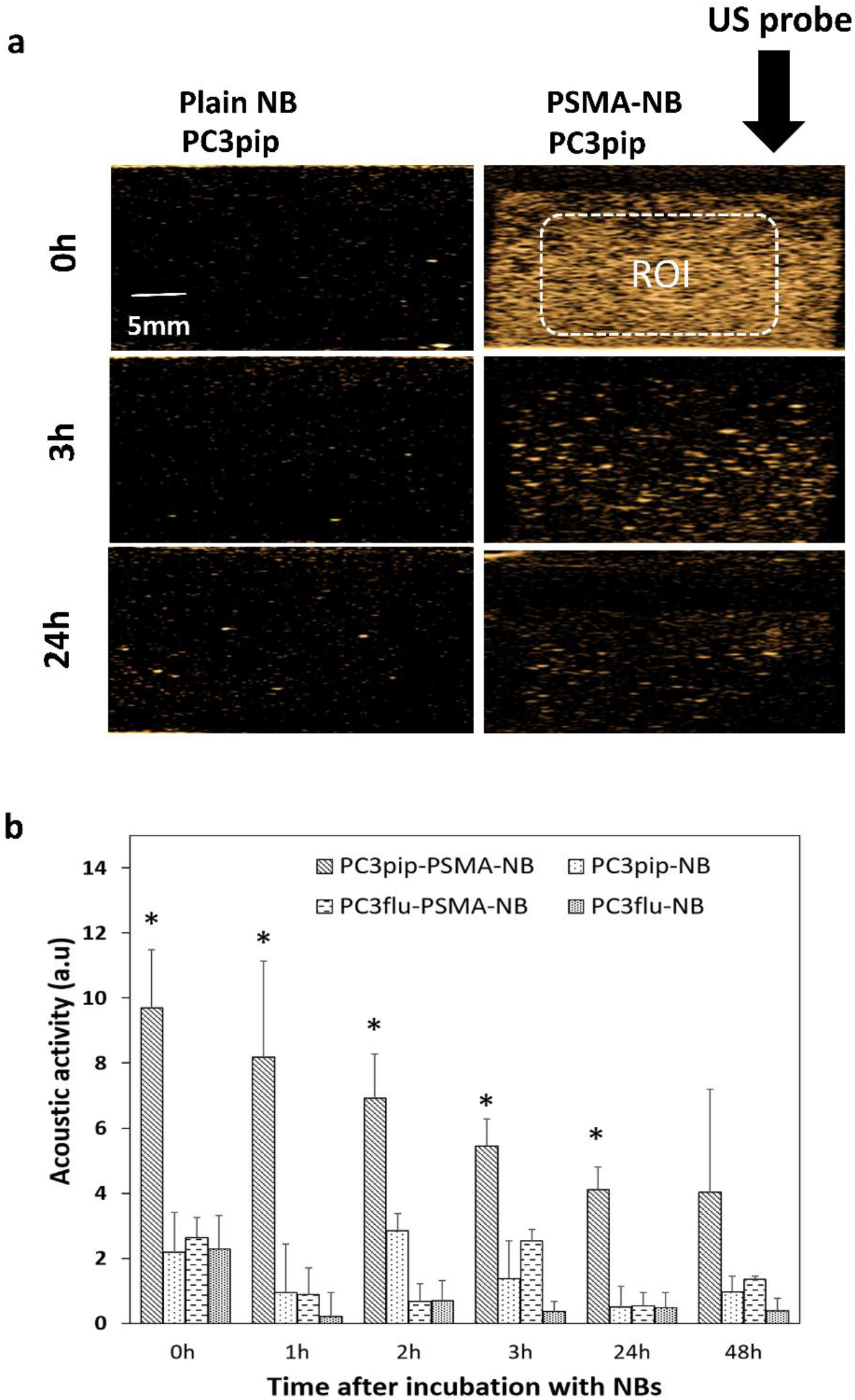
(A) Representative US images of PSMA-positive PC3pip cells incubated with PSMA-NB at different time points after the initial 1 h NB exposure. (B) Acoustic activity of PSMA-NB and NB incubated PC3pip cells and PSMA-negative PC3flu cells at different times post treatment. PSMA-NB incubated PC3pip shows significantly high acoustic activity at t=0 to t=24 h time points compared to all the other groups. n=3, error bars represent mean ± s.d., * denotes statistically significant differences (p<0.05) from all other groups at each time point.

### 2.12 Statistical analysis

Statistical analysis was carried out using Microsoft Excel and Origin. An unpaired Student’s t-test (two-tailed) and one-way ANOVA were used to compare between groups. Data are presented as a mean ± STD (standard-deviation). The experiments were repeated at least three times, unless stated otherwise.

## 3. Results and discussion

### 3.1 Nanobubble characterization

Nanobubble preparation, functionalization with the PSMA-1 ligand and verification of the lipid-ligand conjugation have been reported previously.^[23]^ The diameter of NB and PSMA-NB used in these experiments was characterized by resonant mass measurement (RMM), and was 281 ± 2 nm and 277 ± 11 nm, respectively. Here the standard deviation refers to the reproducibility of mean bubble diameter over three formulations. The range of bubble diameters for each formulation was 111– 692 nm and 110 - 673 nm, respectively. Validation of the RMM analysis and its optimization for use in NB characterization has been previously described^[18, 23]^

The concentration of PSMA-NB (4 × 10^11^ ± 2.45 ×10^10^ NB/ml) was similar to the that of NB (3.9 × 10^11^ ± 2.82 ×10^10^ NB/ml).

### 3.2 Ultrasound signal in tumor rim and core

*In vivo* experiments were performed using the FUJIFILM VisualSonics Vevo 3100 high frequency ultrasound scanner. A total of 9 mice bearing subcutaneous human PC3pip tumor xenografts were used for the experiment. The PC3pip cells were retrovirally transfected to express the PSMA receptor.^[20]^ Animals were divided into 3 groups: those receiving exams with PSMA-NB, NB or Lumason MB. *In vivo* bubble kinetics were imaged with 2D nonlinear contrast mode after injection of 200 μl of undiluted either PSMA-NB (8 × 10^10^ NBs), NB or Lumason via tail vein. Figure 1a shows the timeline for the experiments. A baseline scan was performed with nonlinear contrast mode in 2D and 3D before bubble injection. To obtain wash-in bubble dynamics, each tumor was scanned for ~3 min at 5 frames per second (fps). Figure 1b, c, d and supplementary Figures S1, S2, S3, show representative US images of tumors after receiving an intravenous injection of either PSMA targeted NB, Plain NB or Lumason MB. The tumor rim and core were delineated by drawing regions of interest (ROIs) first on the B-mode images, where the skin and tumor can be clearly separated. The skin was excluded from the analysis. The same ROIs were then applied to the nonlinear image, and the average signal intensity as function of time (time intensity curve-TIC) was calculated for the tumor periphery and tumor core ROIs. Figure 1e shows the TIC curves corresponding to the tumor rim and core obtained with the first 1000 frames at 5 fps (Figure S4 a) followed by a second scan of 1000 frames at 1 fps. Before changing the frame rates from 5 fps to 1 fps, a 3D US scan was performed to observe the bubble distribution in the whole tumor mass.

Rapid signal enhancement was observed in tumors imaged with either PSMA-NB or plain NB, reaching peak intensity in 1 to 2 min post-injection. There was no significant difference in peak enhancement (PE) of entire tumor for PSMA-NB and NB, which was an indication of the similar initial transport of targeted and untargeted bubbles (Figure S4 b). The variability of bubble kinetics within each tumor was further investigated by analyzing the tumor rim and core separately. Here, a loop ROI was drawn around the tumor rim (periphery, carefully separated from the skin), and the tumor core (Figure 1). As shown in Figure 1e, immediately after injection of either PSMA-NB or plain NB, enhancement of the tumor rim was faster than that in tumor core. In contrast to tumor rim wash-in rate, both NB and PSMA-NB demonstrated slower wash-in to the tumor core compared to the tumor rim. Quantitative TIC parameters were calculated and are summarized in Table 1. The time to peak (TTP) in tumor core for the PSMA-NB was significantly longer than that of plain NB (2.48 ± 0.71 min vs 1.21 ± 0.15 min, p×0.05). However, the time to peak (TTP) in tumor rim for PSMA-NB and NB did not show significant differences. Likewise, there was no significant difference in wash-in area under the curve (WiAUC) for both groups. Similar peak enhancement (PE) was also seen in both the tumor rim and core with both NB and PSMA-NB. However, within each NB type, the PE between rim and core was significantly different for both NB group (Table 1). The washout AUC for the tumor rim and tumor core with PSMA-NB was significantly longer than that of the NB group (Figure 1e, Table 1).

**Table 1.** Summary of quantitative kinetic parameters of tumor rim and core obtained from time intensity curve (TIC)

| Bubble Type |  | Peak Enhancement (PE) (a.u) | Time to peak (TTP)(min) | Wash-in AUC (WiAUC) | Washout AUC | Total AUC |
| --- | --- | --- | --- | --- | --- | --- |
| PSMA-NB | Rim | 26263 ± 1930* | 1.77 ± 0.71 | 28718 ± 12677<sup>α</sup> | 187128 ± 49070<sup>#</sup> | 215847 ± 53019<sup>#</sup> |
| NB | Rim | 25099 ± 7109* | 1.52 ± 0.62 | 29354 ± 14462<sup>α</sup> | 89885 ± 30974<sup>#</sup> | 119239 ± 27357<sup>#</sup> |
| Lumason | Rim | 559 ± 118* | 0.84 ± 0.07 | 170 ± 19<sup>α</sup> | 3322 ± 356<sup>#</sup> | 3492 ± 372<sup>#</sup> |
| PSMA-NB | Core | 9175 ± 7871 | 2.48 ± 0.71<sup>#</sup> | 9253 ± 5744 | 45163 ± 23254 | 46591 ± 29509 |
| NB | Core | 7445 ± 1963 | 1.21 ± 0.15<sup>#</sup> | 5142 ± 2306 | 28965 ± 11042 | 29603 ± 9847 |
| Lumason | Core | 157 ± 28 | 0.77 ± 0.11<sup>#</sup> | 59 ± 18 | 1698 ± 158 | 1757 ± 157 |

All values expresses as mean ± s.d., groups with the similar symbols were statistically significant different; \* P < 0.001, #p < 0.01, <sup>α</sup>p < 0.01.

As an additional control, NB accumulation was also compared with the commercially available MB contrast agent Lumason®. The contrast enhancement occurred rapidly with Lumason (TTP is 0.84 ± 0.06 min for rim and 0.77 ± 0.11 for core) and it was significantly different from that of both kinds of NBs. Furthermore, the PE and the WiAUC of Lumason were significantly lower compared to the two NB groups. Importantly, the nonlinear contrast imaging parameters used for bubble studies were kept constant, meaning that the transmit frequency used to image Lumason was higher than typically utilized for this agent. This could result in lower sensitivity of detection and subsequent reduction in signal enhancement. However, relative contrast agent kinetics should be relatively unaffected.

The large observed differences in NB dynamics between the rim and core of each tumor can be attributed to factors related to vascular heterogeneity between the two regions and the PSMA biomarker distribution in the tumor mass ^[24–26] [27]^. Microvessel density (MVD), which is typically greater at the tumor periphery,^[28, 29]^ will increase the rate of signal increase in the region and the likelihood of extravasation. Once within the tumor microenvironment, PSMA targeted NBs are expected to bind to the biomarker, which may delay the rate of transport out of the region and reduce the available concentration in circulation. Hence, the time to reach PE with PSMA-NB is slower compared to the freely flowing NB. The washout AUC for PSMA-NB was also significantly higher than that of NB, which is consistent with high retention of targeted NB in the tumor.

### 3.3 Whole tumor imaging with 3D ultrasound

To gain a better understanding of the bubble distribution in the whole tumor mass, 3D US was implemented after administration of PSMA-NB, NB, and Lumason MB. A slice thickness of 0.05 mm was used to reconstruct the tumor bubble distribution in the entire volume at the baseline, at the peak and at the endpoint of the study. The scans were initially carried out in conjunction with the 2D kinetic imaging (Figure S5). The 3D analysis software provided by the scanner manufacturer calculates the percent of voxels which display nonlinear signal within the analysis volume. In agreement with 2D scanning, the 3D analysis also demonstrated a similar signal at the peak for both PSMA-NB and NB (Figure 2b and c). At 3 min, both PSMA-NB and NB covered about ~90% of the tumor rim (86.9 ± 0.8% and 87.7 ± 6.6 % respectively, *p* = 0.6). Similarly, PSMA-NB and NB were detected in ~60% of the tumor core (64.9 ± 14.5% and 62.4 ± 28.1 % respectively). The percent of agent of Lumason detected in tumor rim was significantly lower compared to the PSMA-NB and NB. At 3 min, Lumason was detected in 10 % (5.7 ± 2.1%) of the tumor rim.

Interestingly, when tumors were imaged with continuous nonlinear ultrasound (1 fps for 16 min after the 3D acquisition), there was no significant difference in nonlinear contrast coverage observed between PSMA-NB and NB groups (Supplementary Figure S6). Also, the ratio of PSMA-NB and NB signal in the whole tumor was lower compared to the previous observation with 2D scanning^[5]^. We speculate that the continuous exposure of NBs which were immobilized within the tumor tissue to the US with may have resulted in rapid bubble dissolution. To test this hypothesis, another set of experiments was carried out, where NBs, PSMA-NBs or Lumason were injected into the mouse, but then were permitted to circulate for 30 min without US exposure. In these experiments, PSMA-NB showed significantly higher percentage of nonlinear signal in the tumor core compared to both NB and Lumason (Figure S5, 25.2 ± 1.5%, 13.9 ± 5.1%, and 0.4 ± 0.4 % respectively, *p* < 0.05) at the experiment end point. According to the 3D tumor analysis, the PSMA-NB also accumulated more in the tumor rim compared to NBs, but the difference was not statistically significant (54.2 ±4.8 %, 38.7 ± 14.4% respectively).

### 3.4 Tumor extravasation studies with 3D ultrasound

To examine intact NB extravasation and accumulation in the entire tumor, mouse perfusion by cardiac puncture was performed at 25 min post-injection and the 3D US scan was performed before and after cardiac puncture (Figure 2a). The cardiac flush procedure eliminates the blood in the vasculature and removes any material including freely moving nanobubbles from the tumor vasculature. After perfusion, the 3D US signal in the tumor corresponds to the remaining materials in the whole tumor parenchyma or intracellular space.

Before perfusion, at 25 min post injection, the PSMA-NB signal in tumor rim was 1.4-fold higher than that of NB (Figure 3b, c; 47.8 ± 19.75 vs 33.2 ± 20.04%, *p* = 0.5). The amount of PSMA-NB in the tumor core was 2-fold higher than that of NB (37.7 ± 20.1% vs 18.9 ± 18.7%). After perfusion, the US signal was reduced in both groups. The US signal in tumor rim for PSMA-NB and NB reduced by 67% and 92%, respectively, relative to the peak values (16.0 ± 8.3% vs 1.5 ± 1.8 % respectively).Furthermore after perfusion, the NB signal was significantly reduced in the tumor core compared to the PSMA-NB (12.2 ± 2.3 % vs 3.2 ± 2.2% respectively, p < 0.05).). The significantly higher (~4-fold) 3D US signal for PSMA-NB in tumor core compared to NBs suggests higher extravasation and retention of PSMA-NB in the tumor core environment. The presence of US contrast after cardiac perfusion provided evidence that intact NB accumulation and extravasation in the tumor parenchyma. As expected Lumason percentage was negligibly small at 25 min post injection (2.0 ± 1.0 for the rim and 0.2 ± 1.2 for the core) and after perfusion (1.3 ± 0.2 for the rim and 0.4 ± 1.0 for the core) in both tumor rim and core.

### 3.6 Histology analysis

Histological analysis was performed after the injection of Cy5.5-PSMA-NB or Cy5.5-NB into tumor tissues. PSMA expression, CD31 expression, and the PSMA-NB and NB distribution were evaluated in each tumor rim and tumor core. The bubbles were labeled with a fluorescent dye; Cy5.5, before the injection. As described in the extravasation studies, 25 min after the bubbles were injected, a 3D US scan was performed to collect the nonlinear signal data, and the animal was perfused with PBS by cardiac puncture. A repeat 3D US scan was performed after perfusion and the tumor was then harvested for histological analysis. Tumor core and rim were imaged and analyzed separately. CD31 expression was, on average, higher in the tumor rim compared to the tumor core (Figure 3a, b). This is consistent with typical tumor physiology, where the periphery tends to be highly vascularized compared to the core.^[30]^ PSMA expression was similar in the tumor periphery and the tumor core (Figure 3a,b). The Cy5.5-PSMA-NB signal was distributed more evenly in the tumor providing additional evidence that PSMA-targeted NB can robustly extravasate from the vasculature and accumulate in the tumor extracellular matrix (Figure 3a). Quantification of histology signal reveal that the Cy5.5-PSMA-NB signal in both tumor core and the rim was significantly higher (3-fold) compared to that of plain NB (*p* < 0.001) (Figure 4c). Enhanced interaction of contrast agent and tumor cells accounts for the high accumulation of PSMA-NB after extravasation.

### 3.7 Confocal imaging of the PSMA-NB internalization in PC3pip cells

PSMA functions as a cell membrane receptor and internalizes the PSMA targeting ligands along with the payload that is attached to the targeting agents^[31]^. When the PSMA ligand binds to the biomarker on the cell membrane, the cell membrane invaginates and the entire particle is engulfed by the cell^[31]^. To investigate the localization of internalized PSMA-NB within the cells, confocal microscopic studies were performed using a fluorescence dye, Lysotracker Red, which stains late endosomes and lysosomal structures.

Our previous fluorescence imaging data showed that the PSMA targeted NB are selectively internalized by the PC3pip cells^[5]^. Confocal imaging results here show internalization, and more specifically, receptor-mediated endocytosis of PSMA-NBs by PC3pip cells (Figure 4a, 100X, Figure S7, 40X). Substantial co-localization of Lysotracker Red (green) and Rhodamine-labeled PSMA-NBs (red) was noted in all PSMA-expressing cells. As shown in Figure 4a and Figure S7, plain NB showed nonspecific uptake by PC3pip cells but with limited late endosomal/ lysosomal co-localization. PSMA-NB showed significantly higher uptake and higher degree of co-localization of PSMA-NB with the late endosomal/ lysosomal vesicles than untargeted NBs in PC3pip cells (Figure 4a).

Imaging of PC3pip cells at 24 h after bubble exposure revealed a lower fluorescence signal compared to the earlier time points (Figure 4b). However, these images also showed a high degree of co-localization of PSMA-NB in late endosomal/lysosomal vesicles. Very low or no fluorescence signal was observed in the cytoplasm or late endosomal/lysosomal compartments of cells incubated with untargeted NB after 24 h. Furthermore, the images showed yellow color staining near the nucleus, which suggested that the majority of the PSMA-NBs were co-localized with either later endosomes or the lysosomes and transported into cell cytoplasm. A higher amount of Lysotracker staining was observed when PC3pip cells were incubated with targeted NBs compared to untargeted NBs, which correlates with mechanism of internalization being receptor mediated and entering into the late endosome/lysosome pathway. PSMA has a unique internalization motif^[32]^ and is reported to have a robust baseline internalization rate of 60 % of its surface PSMA in 2 h. Transmembrane location and internalization make PSMA an ideal target for imaging and therapy^[6, 20, 33–36]^. Overall, PSMA mediated endocytosis appears to be the main pathway for the internalization of PSMA-targeted nanobubbles in PSMA-expressing prostate cancer PC3pip cells.

### 3.8 Persistence of PSMA-NB *in vitro*

To investigate the effect of cellular internalization of nanobubbles on the persistence of acoustic activity, we compared effects of passive cellular uptake versus receptor-mediated endocytosis of PSMA-targeted NBs in prostate cancer cells. We first investigated the kinetics of nonlinear acoustic properties of both PSMA-positive PC3pip and PSMA-negative PC3flu cells after incubation for 1 h with PSMA targeted or untargeted NBs by harvesting cells and imaging them with the same CHI imaging pulse parameters as used for the mouse studies. As shown in Figure 5, after incubation for 1h, PC3pip cells incubated with PSMA-NB showed 3.25 fold higher acoustic activity compared to the plain NB incubated PC3pip cells (Figure 5; p < 0.05; 9.69 ± 1.78 dB vs, 2.19 ± 1.22 dB respectively) and showed higher signal intensity compared to all other groups (NB in PC3pip cells, PSMA-NB in PC3flu cells and plain NB in PC3flu cells) and the negative control (cells only). Furthermore, the significantly higher acoustic activity for internalized PSMA-NBs persisted for all the time points tested except at 48h. (Figure 5 and Figure S8). After 24 h, PC3pip cells exposed to PSMA-NBs showed significantly higher US signal intensity (4.11 ± 0.68 dB; p< 0.05) compared to all other groups. PSMA-NBs incubated under the same conditions but without cells, showed an initial signal intensity comparable to the PSMA-NB incubated with PC3pip cells (Figure S9). However, after the 3h time point, the signal intensity of PSMA-NB without cells was significantly lower than compared to the PSMA-NB internalized in PC3pip cells, indicating of the extended presence of gas in the cellular environment compared to the free PSMA-NB in solution.

It has been previously shown that either internalized or membrane-bound MB are protected by much greater viscous damping by cells^19^ compared to free MB. In agreement with these findings, we also observed that NBs internalized into cells as visualized by the confocal images showed significantly higher backscatter for a longer period of time compared to the free NB under the same conditions (Figure 5, Figure S8, S9). Furthermore, at later time points, PSMA-NB in PC3pip cells showed higher contrast than the plain NB incubated with either PC3pip or PC3flu cells and PSMA-NB incubated with the PSMA-negative PC3flu cells. Nonspecific uptake of NB by the cells slightly decreases the rate of signal decay from the NBs. However, a much slower decay was observed for the PSMA-NBs localized within the endosomes. Thus, we postulated that the long-time survival of PSMA-NB in cells might be due to stabilization by the endosomal/lysosomal entrapment.

### 3.9 Analysis of C3F8 in cells using headspace GC/MS

To validate that intact, gas-bearing nanobubbles internalized into and remain in the PC3pip cells, cells were harvested 3 h following exposure to PSMA-NB or NB, and the presence of C_3_F_8_ inside the cells was analyzed using headspace GC/MS. The use of headspace GC/MS to quantify C_3_F_8_ gas concertation in NBs was previously reported and validated^[18]^. The analysis was performed using the relative abundance of the peak observed at the mass to charge ratio (m/z) of 169, which is a characteristic peak for C_3_F_8_. NBs at different concentrations were used to generate the calibration plot with GC/MS (Figure 6a and b). We observed a linear relationship between peak area (the area under the curve of abundance vs time curve) and the number of bubbles. The GC/MS data showed that the peak area obtained from the PSMA-NB incubated PC3pip cells showed a 3.5-fold higher value compared to that of plain NB incubated PC3pip cell suspension (Figure 6c; Peak area of PSMA-NB, NB, and cells were 16778 (a.u), 6274 (a.u), and 2172 (a.u) respectively). Based on the calibration curve, this corresponds to approximately 500 average-sized PSMA-NB versus 138 average-size plain NB per cell. To the best of our knowledge, this is the most direct method for confirming the presence of gas vesicles within the cells. The ratio of the peak area is consistent with the difference in acoustic activity from the cells as shown in Figure 6. It is also strikingly similar to the difference in the ultrasound signal seen from PSMA-NB versus plain NB accumulation in PC3pip tumors after clearance of circulating NBs^[5]^ and supports the data showing that *in vivo* the targeted NB extravasated and were retained in the tumors in an intact form^[4]^.

**Figure 6.**
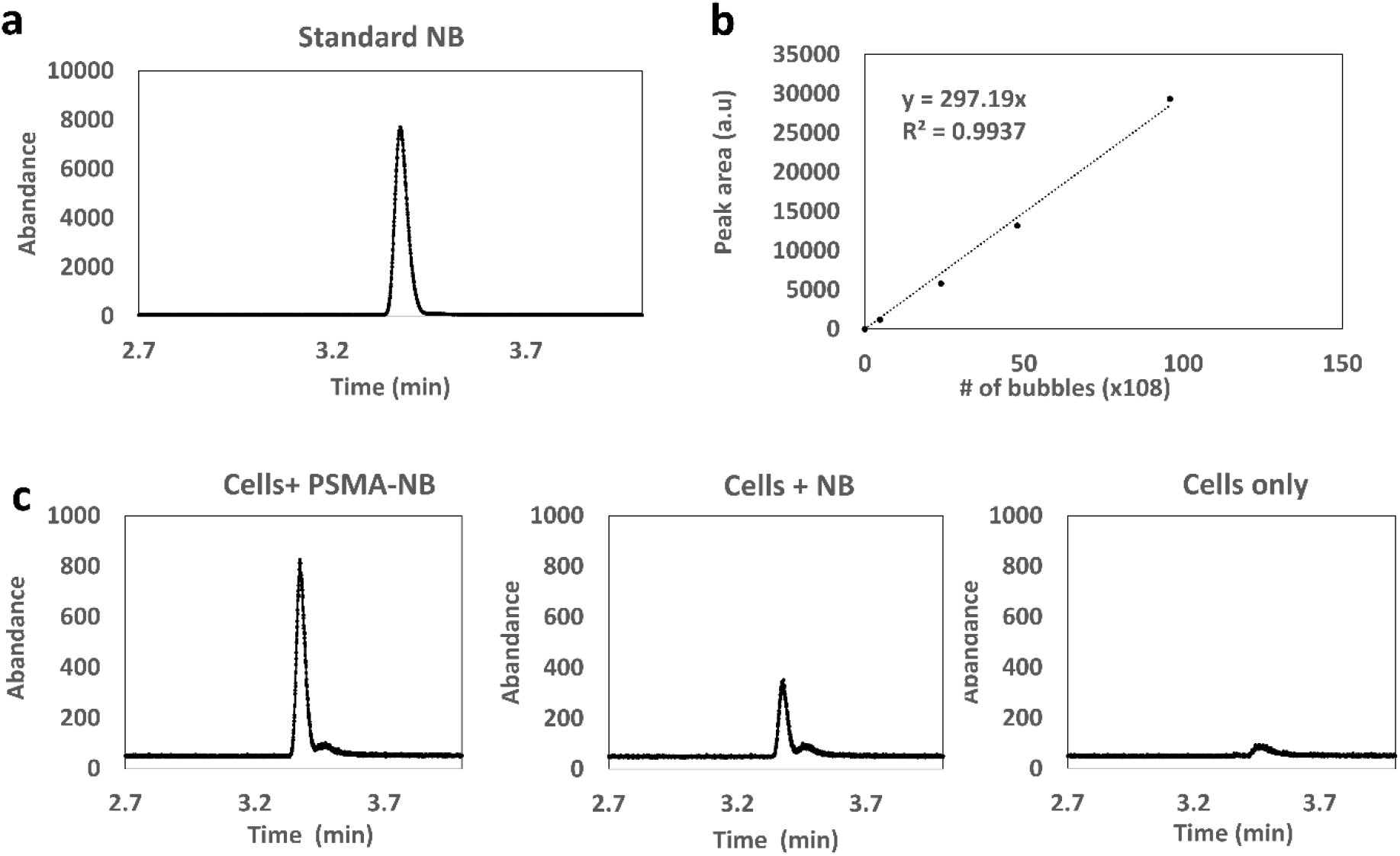
(A) Head-space GC/MS analysis of C_3_F_8_ gas generated by NB; eluting at 3.37min (B) Calibration curve for various concentrations of bubbles vs peak area correspond to C_3_F_8_ gas (B) Head space GC/MS analysis of C_3_F_8_ gas generated by PSMA-NB and plain NB internalized PC3pip cell suspension.

### 3.10 *In vivo* application of PSMA-NB internalized cells

Finally, to confirm that the prolonged intracellular retention can also be visualized *in vivo,* we studied the acoustic activity of internalized bubbles in cells upon injection into mice. PC3pip cells incubated with PSMA-NBs were injected subcutaneously into flank area of nude mice and imaged at 12 MHz As a control, cells without exposure to NBs were injected adjacent to the labeled cells injected area. Figure 7 a shows the US images obtained 0-24 h and 8 days after cell injection. PSMA-NB-incubated cells demonstrated significantly high contrast compared to the unlabeled cells immediately after injection (9.7 ± 2.9 vs 5.2 ± 1.5: p < 0.05). The initial signal seen from the control cells is most likely a result of air bubbles entrapped in Matrigel. After 3 and 24 h, the contrast was still significantly higher in regions with PSMA-NB-incubated cells compared to control cells (Figure 7b).

**Figure 7.**
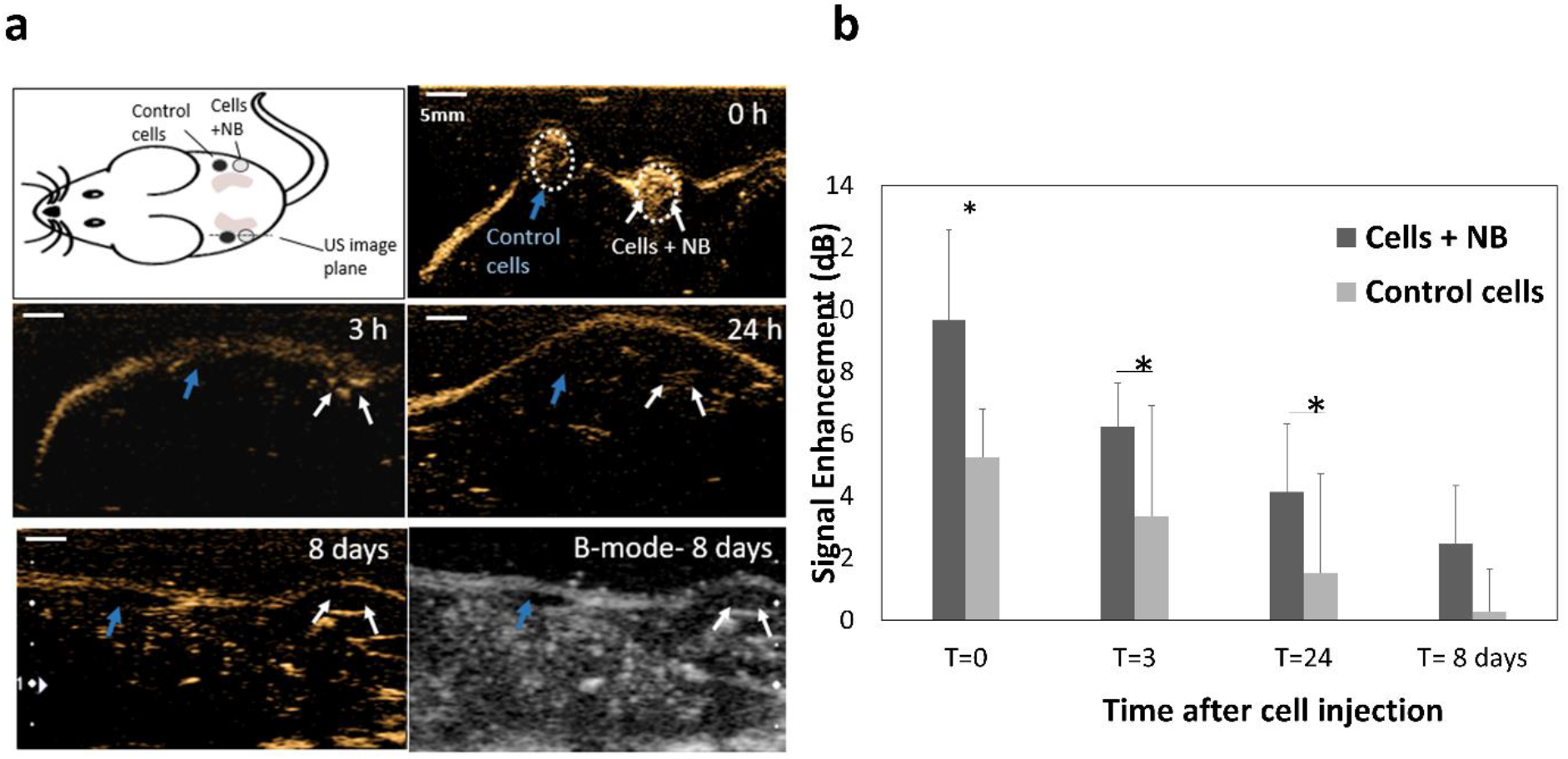
(A) Representative US images of subcutaneous injection of cells incubated with PSMA-NB compared to control unlabeled cells at different time points. Cells were mixed with Matrigel prior to injection. (B) Average US signal intensities at each time points. *p<0.05.

Internalized PSMA-NB in PC3pip cells were primarily co-localized with intracellular vesicles and showed substantial backscatter activity for 48 hours after incubation *in vitro*, and for one week *in vivo*. To the best of our knowledge, this study shows, for the first time, direct evidence of PSMA targeted-NB uptake and extended retention in cancer cells, and demonstrates the significant role of endosomes/ lysosomes in stabilization of the NB acoustic activity. Our findings are consistent with published reports that showed internalized MB are able to enter into or be phagocytosed by cells^[37]^, and some studies demonstrated extended acoustic activity after MB internalized in neuroprogenitor cells^[16]^. However, additional studies are needed to fully understand the intracellular trafficking of nanoscale ultrasound contrast agents. Nonetheless, the stabilized intracellular acoustic activity holds promise for development of specific ultrasound molecular imaging techniques as well as highly targeted and low systemic toxicity therapeutic applications. Shen et. al. demonstrated folate tagged NB selectively accumulated in folate positive endothelial cells and cervical and lung cancer cells via clathrin and caveolin endocytosis^[38]^ which lead to selective cytotoxicity when combined with therapeutic US. In line with these results, the current studies may also open new avenues to focused prostate cancer therapy.

## 4. Conclusion

This study demonstrates that the active targeting of NB to PSMA increases the extravasation and the accumulation of these agents in PSMA expressing tumors. The signal from the PSMA-NBs can be clearly visualized in both 2D and 3 D nonlinear contrast imaging mode, and the prolonged signal enhancement can be consistently visualized throughout the entire tumor volume. Quantification of the dynamic nanobubble-enhanced ultrasound parameters shows that both wash-in and of PSMA-NB are thought to be due to biomarker interaction and binding, as well as cellular internalization. Furthermore, in vitro studies suggested the active targeting of NB to PSMA selectively enhances cellular internalization in PSMA-positive PC3pip cells, and internalized PSMA-NB showed prolonged stability in the cellular environment, most likely due to entrapment in endosomal vesicles. GC/MS analysis further confirms that intact NB persists in cells after internalization. The results presented here support prior studies showing prolonged acoustic activity of targeted nanobubbles in biomarker expressing tumors and open doors for new molecular imaging and targeted therapy approaches using ultrasound.

## Supporting Information

Supporting Information is available.

## Acknowledgements

We would like to acknowledge Al de Leon, Rich Lee from Microscopy Imaging Core, Case Western Reserve University, and Melissa Yin from FUJIFILM VisualSonics, Toronto, Canada for the support given for this project. This work was funded by the National Institutes of Health (R01EB028144), 1S10OD021635-01, and the CWRU Coulter Translational Research Partnership. We also acknowledge additional support from the Case Comprehensive Cancer Center P30CA043703 in the form of a pilot grant and the National Foundation for Cancer Research (NFCR). Views and opinions of, and endorsements by the author(s) do not reflect those of the funding agencies. The authors report no conflicts of interest to this work.

