## Supporting Information for "Intracellular Vesicle Entrapment of Nanobubble Ultrasound Contrast Agents Targeted to PSMA Promotes Prolonged Enhancement and Stability *In Vivo* and *In Vitro*"

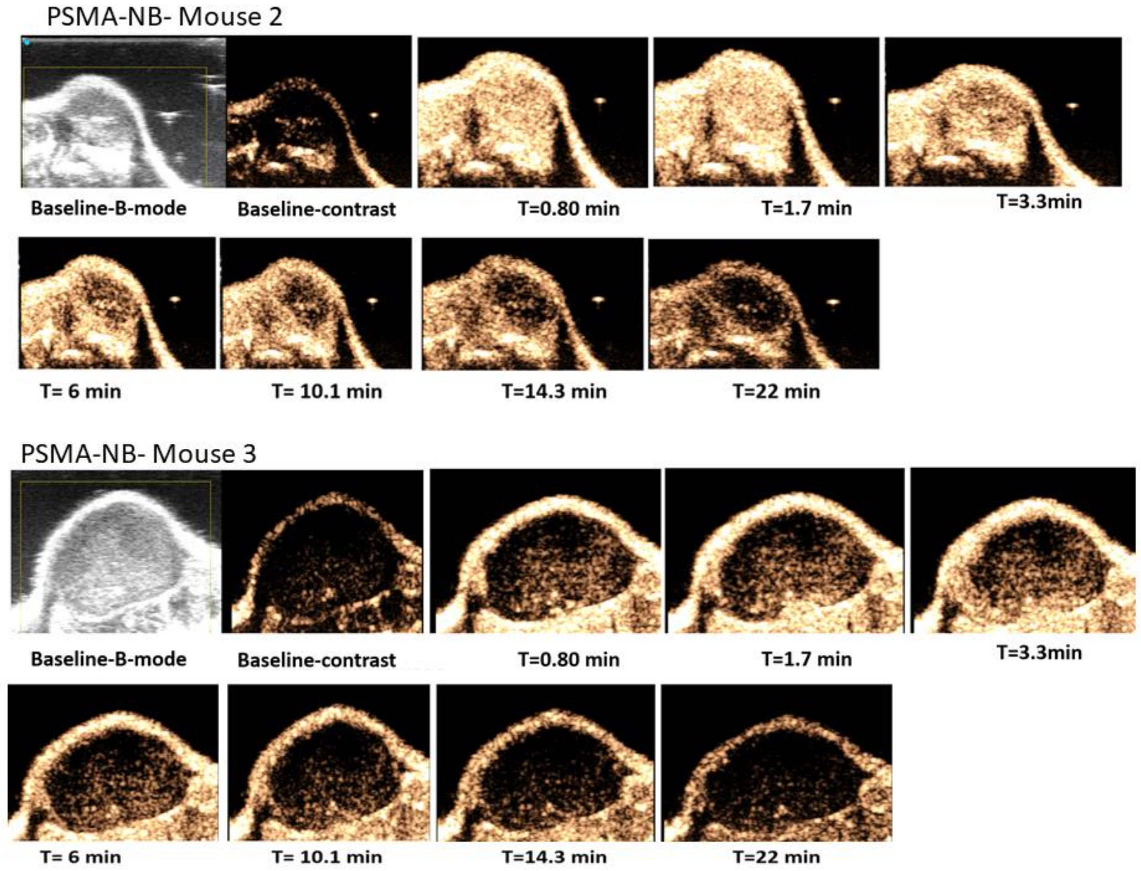


**Figure S1.** Nonlinear contrast enhanced US images showing **PSMA-NB** distribution in tumors of two additional mice at different times after bubble injection.

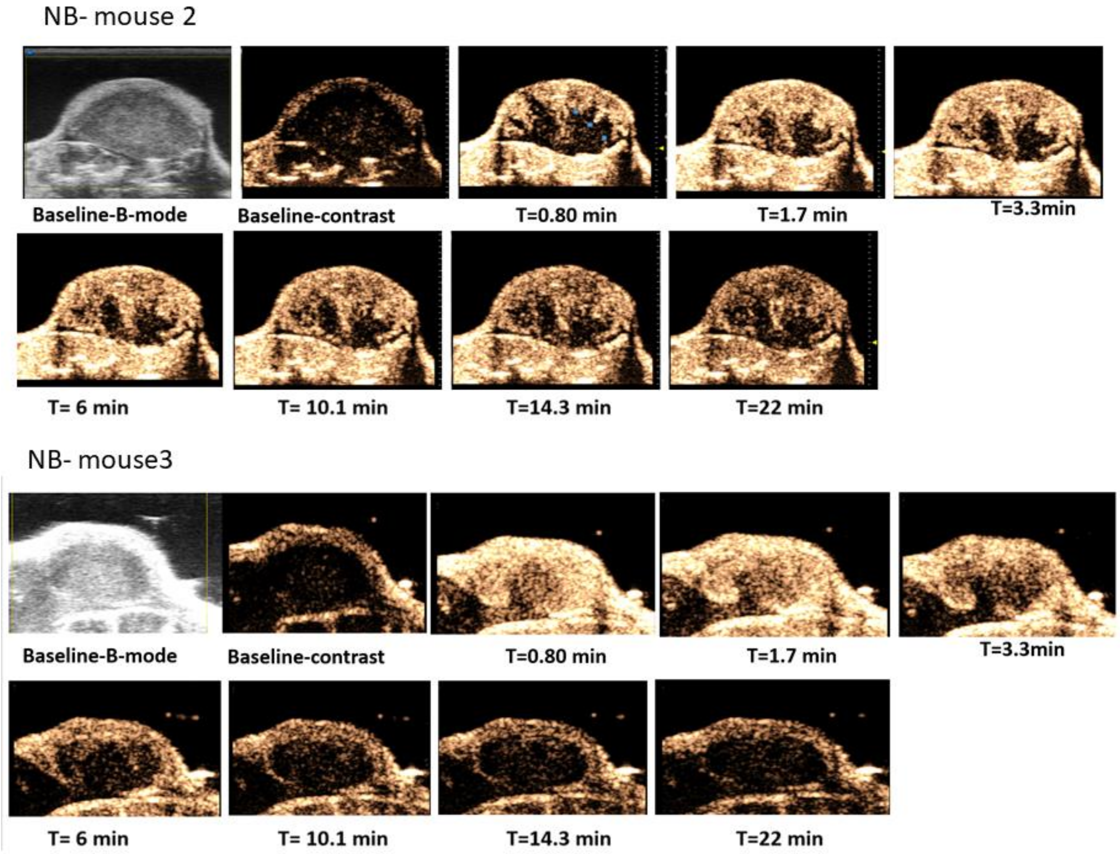


**Figure S2.** Nonlinear contrast enhanced US images showing **NB** distribution in tumors of two additional mice at different times after bubble injection.

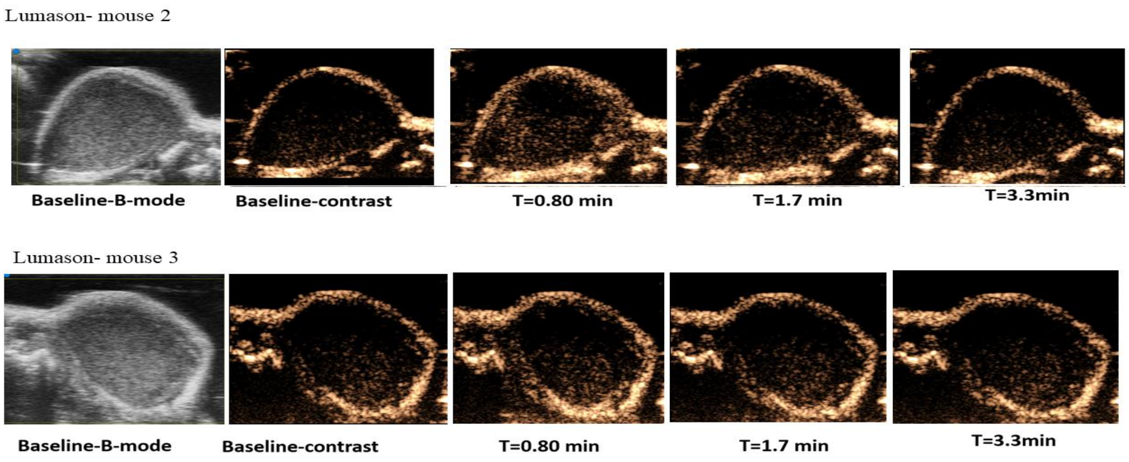


**Figure S3.** Nonlinear contrast enhanced US images showing **Lumason MB** distribution in tumors of two additional mice at different times after bubble injection.

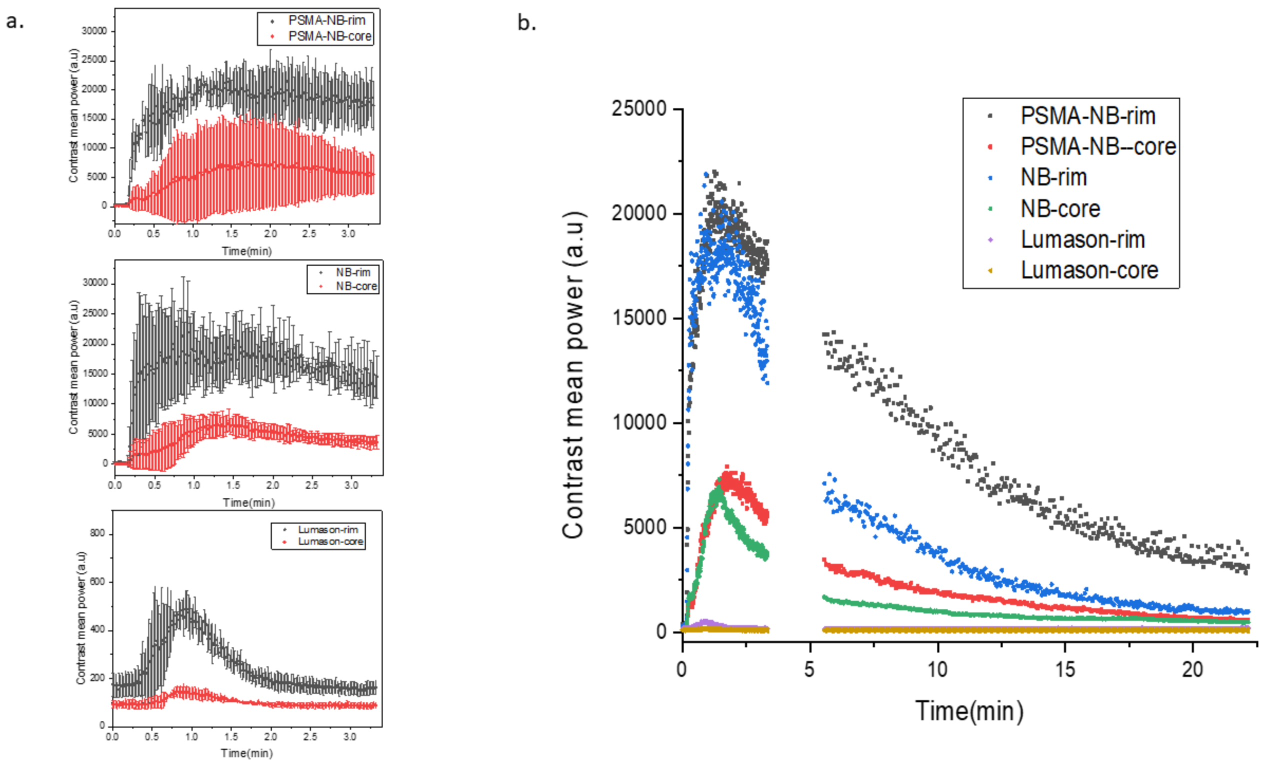


**Figure S4.** a) Average signal intensity for wash-in phase in tumor rim and core as a function of time for PSMA-NB, plain NB, and Lumason MB. b) Average time-intensity curves for both wash-in and washout phase for PSMA-NB, plain NB, and Lumason MB in tumors (without error bars). Tumors were imaged at 18 MHz, 5 frames/second for 3.33 min and 1 frame/second for 16.67 min.

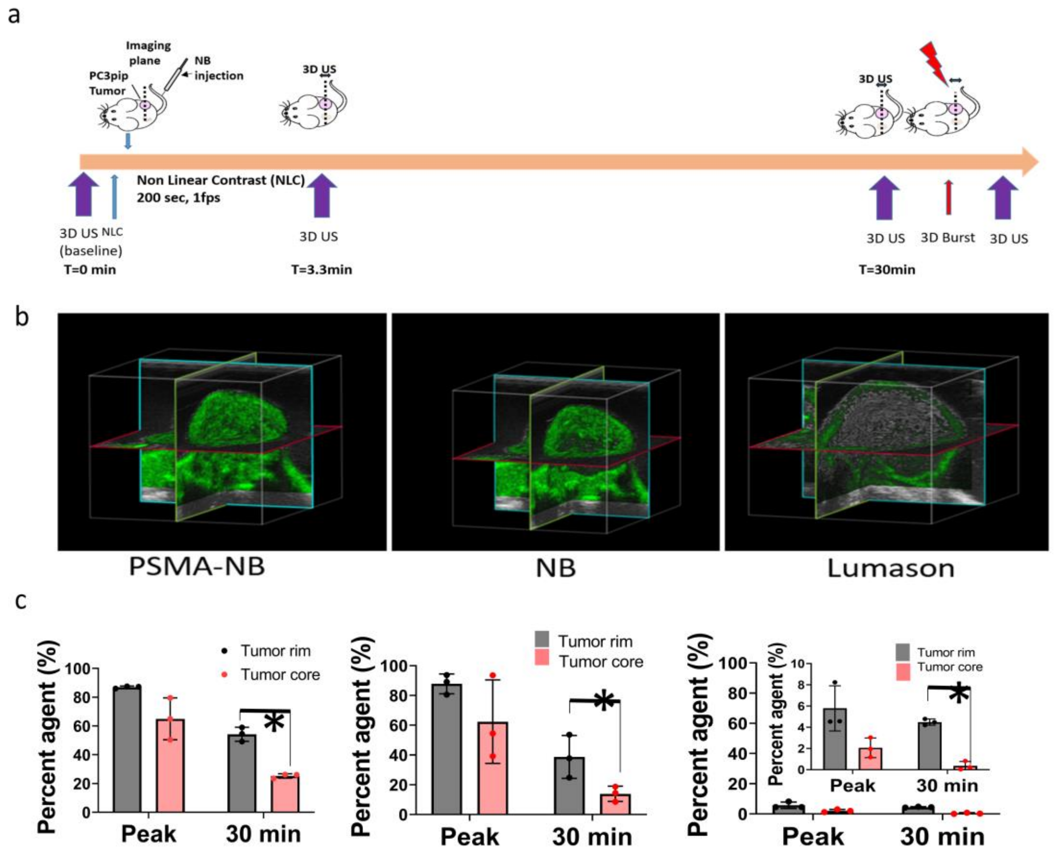


**Figure S5.** 3D ultrasound scanning to visualize bubble distribution in the entire tumor volume a) Timeline showing the 3D US scanning points. b) Representative 3D US images of the tumor showing PSMA-NB, NB, and Lumason at the peak signal intensity. c) Percent agent coverage per tumor type quantified at peak and the t=30 min after baseline subtraction. n=3, error bars represent mean ± s.d., * $p<0.05$.

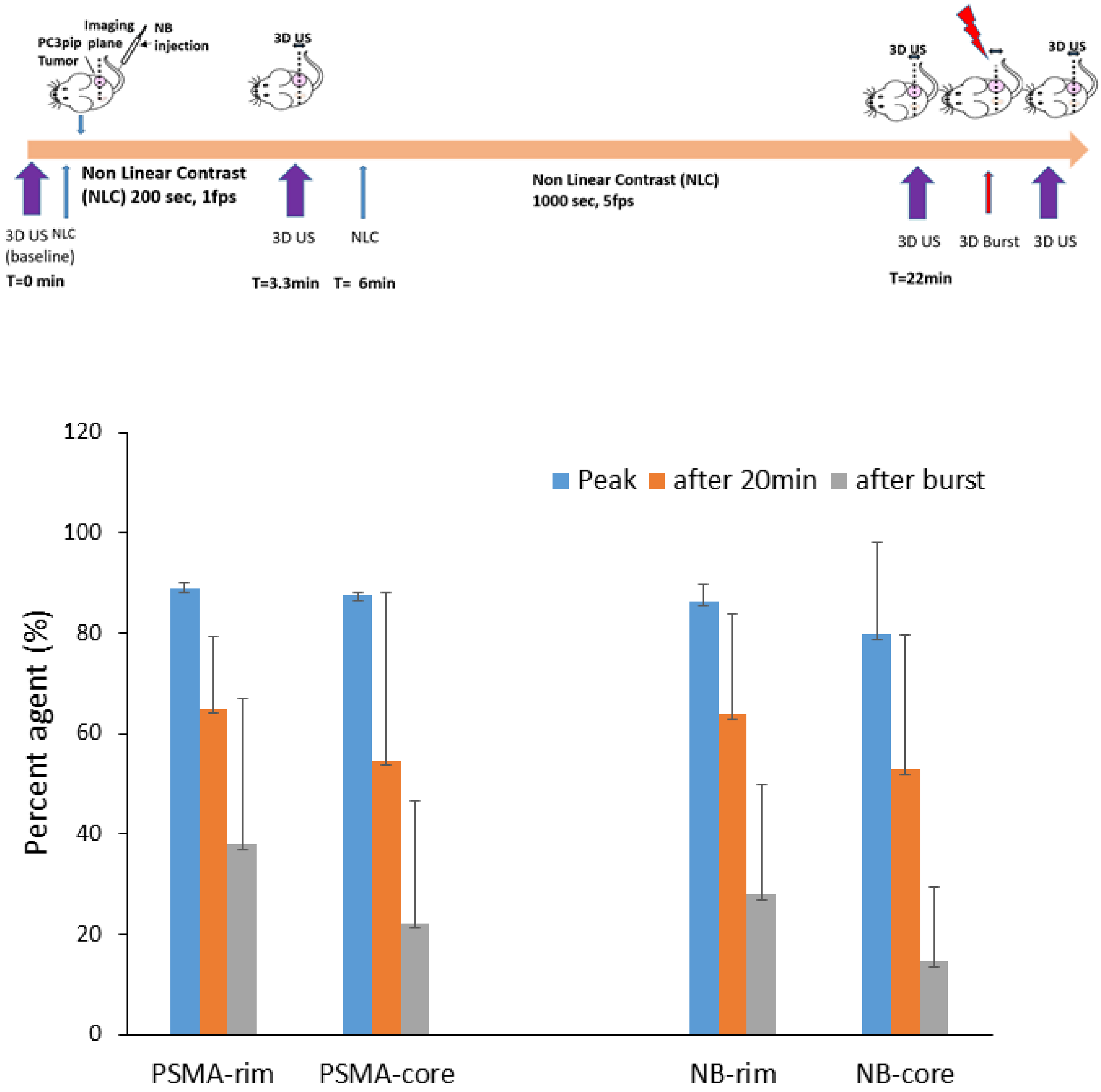


**Figure S6.** a) Timeline showing the 3D US scanning points. b) Quantification of 3D US signal intensities at peak, at t=22 min, and after burst. The values obtained after subtraction of the baseline value.

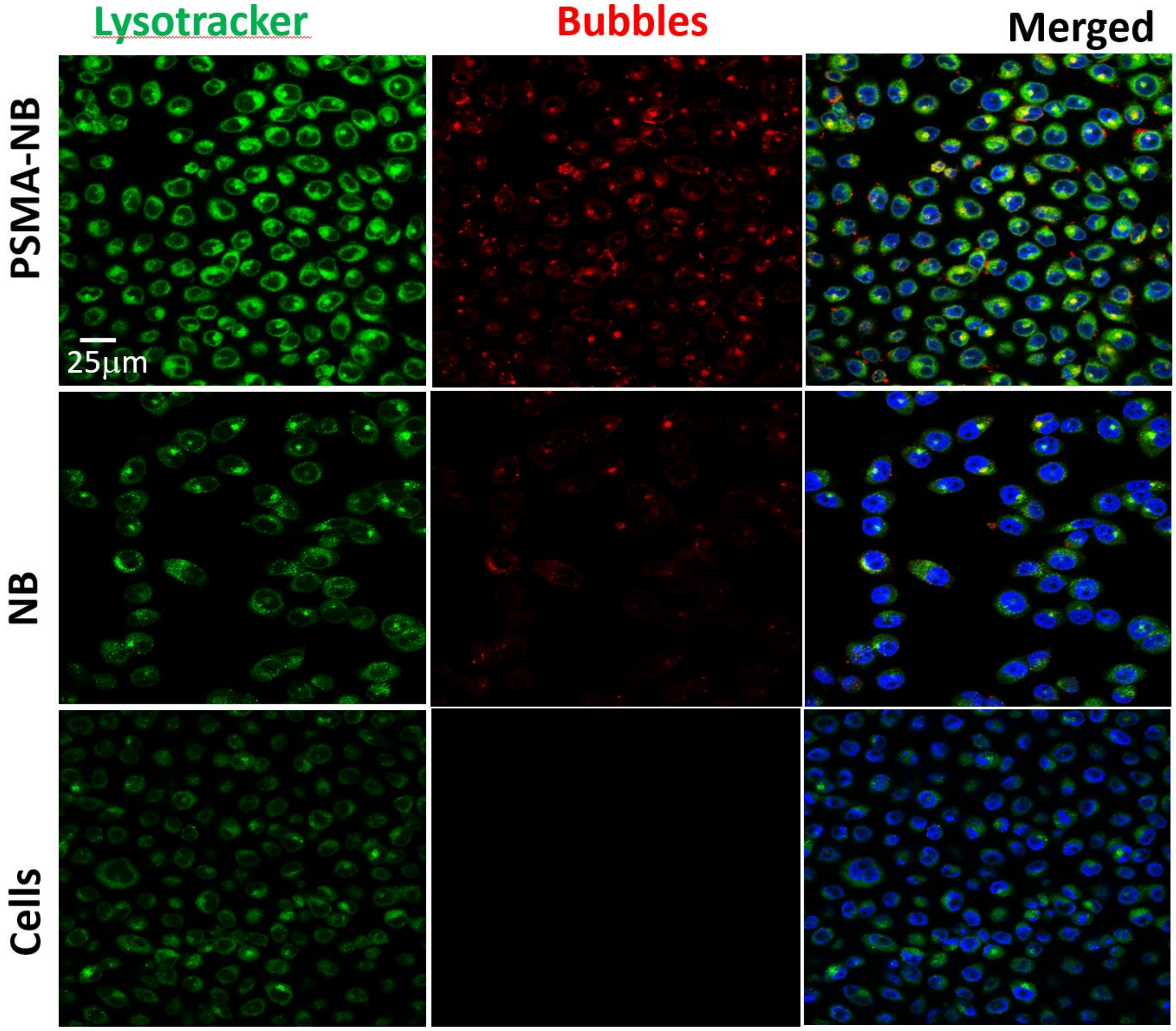


**Figure S7.** Representative confocal images of PSMA-NB and NB distribution in PC3pip cells after 1h incubation; 40X (blue-nuclei, red-NB, and green-late endosome/lysosomes).

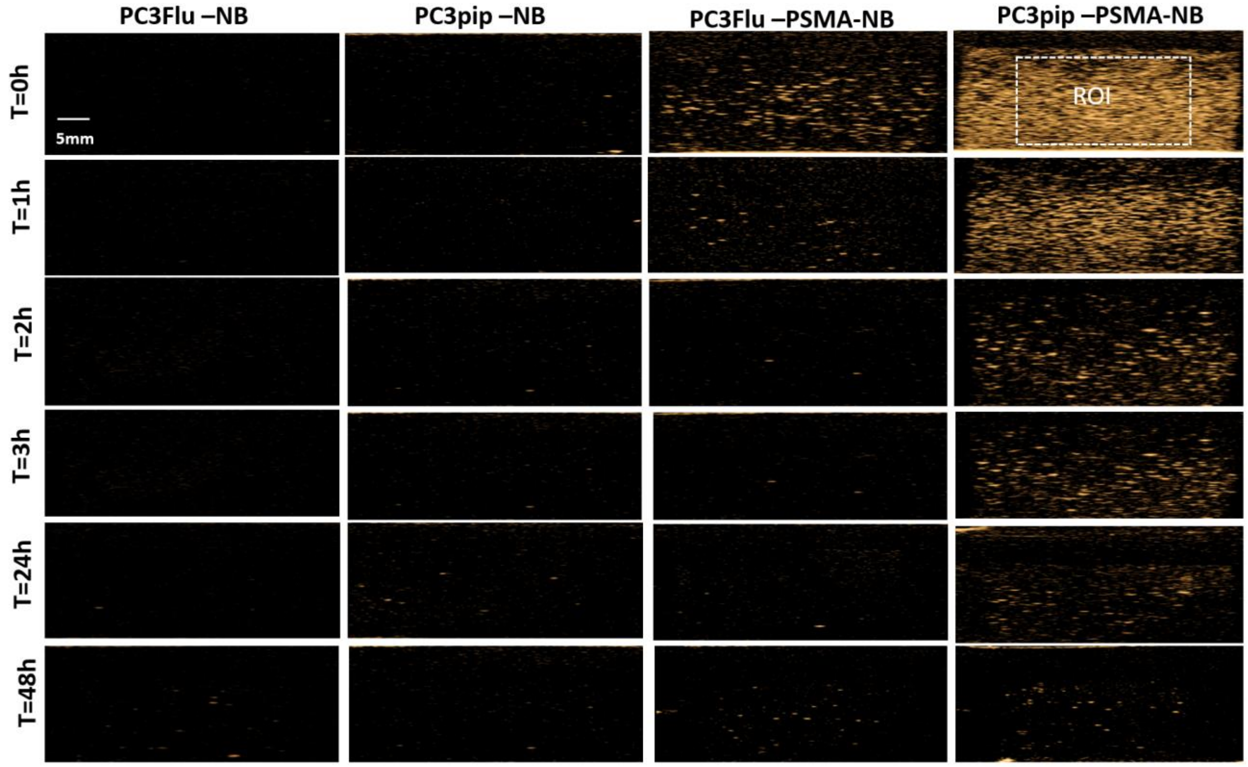


**Figure S8.** Representative US images of PSMA-NB, NB treated PSMA positive PC3pip cells and PSMA negative PC3flu cells at different times post treatment. PSMA-NB incubated PC3pip shows significantly high acoustic activity at t=0 to t=24 h time points compared to all the other groups. n=3.

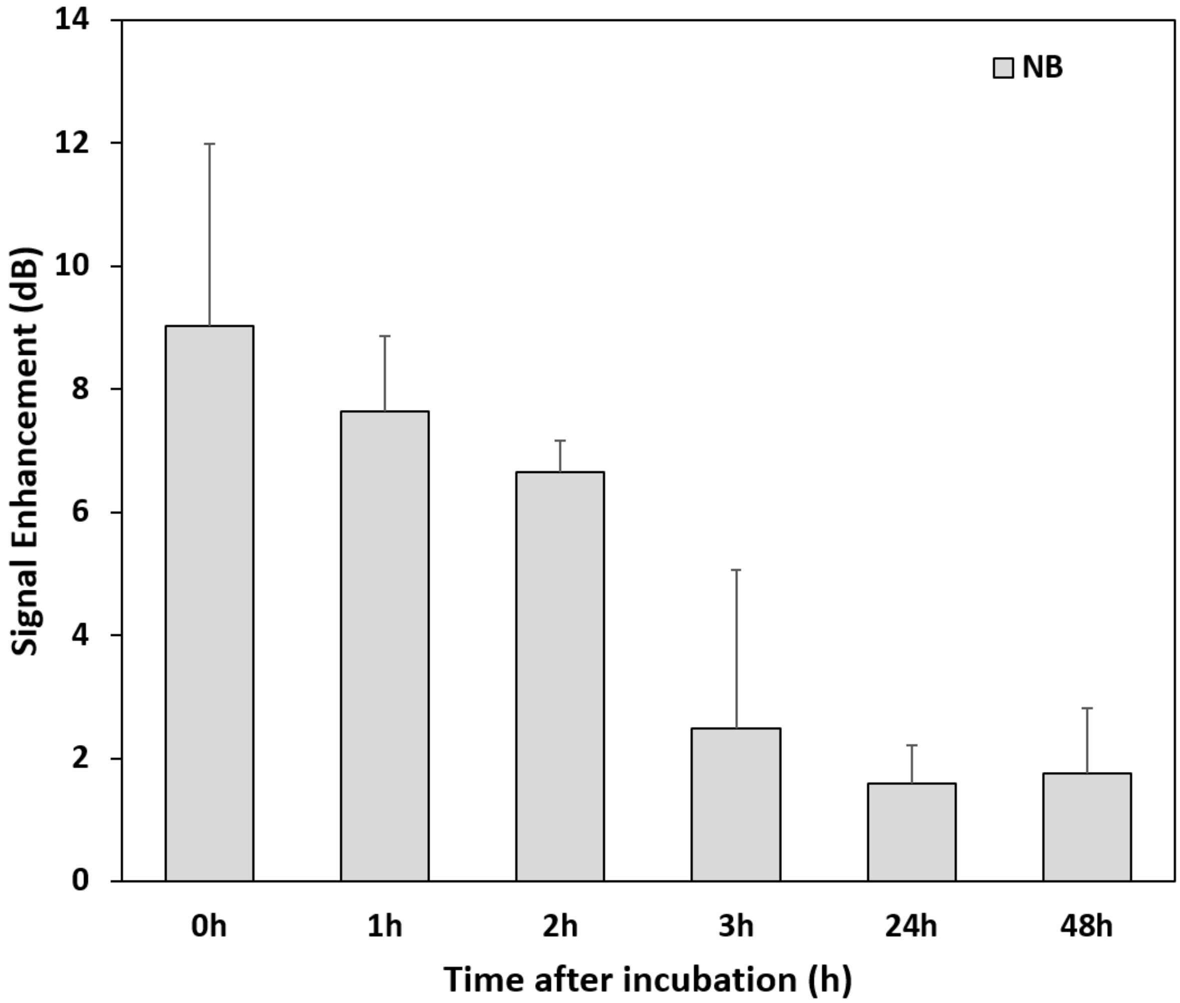


**Figure S9**. Signal enhancement of PSMA-NB incubated under the same conditions as PSMA-NB internalized cells (at 37 °C and 5 % $CO_2$) and imaged at different times points.
